# High spatial resolution global ocean metagenomes from Bio-GO-SHIP repeat hydrography transects

**DOI:** 10.1101/2020.09.06.285056

**Authors:** Alyse A. Larkin, Catherine A. Garcia, Melissa L. Brock, Jenna A. Lee, Nathan Garcia, Lucas J. Ustick, Leticia Barbero, Brendan R. Carter, Rolf E. Sonnerup, Lynne Talley, Glen A. Tarran, Denis L. Volkov, Adam C. Martiny

## Abstract

Detailed descriptions of microbial communities have lagged far behind physical and chemical measurements in the marine environment. Here, we present 720 globally distributed surface ocean metagenomes collected at high spatio-temporal resolution. Our low-cost metagenomic sequencing protocol produced 2.75 terabases of data, where the median number of base pairs per sample was 3.48 billion. The median distance between sampling stations was 26 km. The metagenomic libraries described here were collected as a part of a biological initiative for the Global Ocean Ship-based Hydrographic Investigations Program, or “Bio-GO-SHIP.” One of the primary aims of GO-SHIP is to produce high spatial and vertical resolution measurements of key state variables to directly quantify climate change impacts on ocean environments. By similarly collecting marine metagenomes at high spatiotemporal resolution, we expect that this dataset will help answer questions about the link between microbial communities and biogeochemical fluxes in a changing ocean.

## Background & Summary

A growing list of coordinated scientific efforts have produced deep metagenomic libraries of the surface ocean. Projects such as the Global Ocean Survey, Tara Oceans, and bioGEOTRACES^1–3^ have significantly advanced our understanding of marine microbial biogeography and biodiversity. However, this ever-increasing abundance of metagenomic data raises the question of how do we move beyond analyses of biodiversity to linking microbial traits with ecosystem function and elemental fluxes^4^. In oceanography, it has been widely acknowledged that sparse sampling results in high noise and error rates that in turn prevent the characterization of dynamic chemical balances and limit biogeochemical models^5^. Thus, we propose that an increased emphasis on high resolution spatiotemporal sampling of marine microbial communities would allow for a more mechanistic understanding of the relationship between microbes and ocean biogeochemistry.

The Global Ocean Ship-based Hydrographic Investigations Program (GO-SHIP) seeks to produce high spatial and vertical resolution measurements of physical, chemical, and biological parameters over the full water column. This internationally-organized program coordinates a network of sustained hydrographic sections that are repeatedly measured on an approximately decadal time scale. Compared to autonomous programs such as Argo, which has significantly increased the spatial and temporal resolution of ocean observations^6^, ship-based programs have the advantage of a much broader range of biogeochemical measurement capabilities. To date, repeat hydrography programs have largely focused on physical (light, currents, water column thermohaline structure, etc.) and chemical (nutrients, oxygen, dissolved organic and inorganic carbon, pH, etc.) state variables. This work has significantly improved our understanding of the response of oxygen^7^, pH^8^, calcium carbonate saturation depth^9^, and sea level rise^10^ to global warming and anthropogenic carbon accumulation^11^. By comparison, systematic and sustained biological measurements of the microbial component of ocean ecosystems has lagged far behind.

Here, we present a dataset of 720 ocean surface water metagenomes collected at high spatiotemporal resolution in an effort to more mechanistically link marine microbial traits and biodiversity to both chemical and hydrodynamic ecosystem fluxes as a part of a novel Bio-GO-SHIP sampling program. Samples were collected in the Atlantic, Pacific, and Indian Ocean basins (Fig 1, Table 1). This effort has been supported by GO-SHIP, the Plymouth Marine Laboratory Atlantic Meridional Transect (PML AMT), and three National Science Foundation (NSF) Dimensions of Biodiversity funded cruises (AE1319, BVAL46, and NH1418) (Table 2). Whereas the median distance between Tara Oceans sampling stations was 709 km and the median distance between bioGEOTRACES sampling stations was 191 km, the median distance between sampling stations in the current Bio-GO-SHIP dataset is 26 km (Fig 2). In addition, the majority of Bio-GO-SHIP samples were collected every 4-6 hours, allowing for analysis of diel fluctuations in microbial composition and gene content^12^. We anticipate that our high-resolution sampling scheme will allow for a more detailed examination of the relationship between the broad range of geochemical parameters measured across the various cruises (Table 2) and microbial diversity and traits.

**Figure 1:**
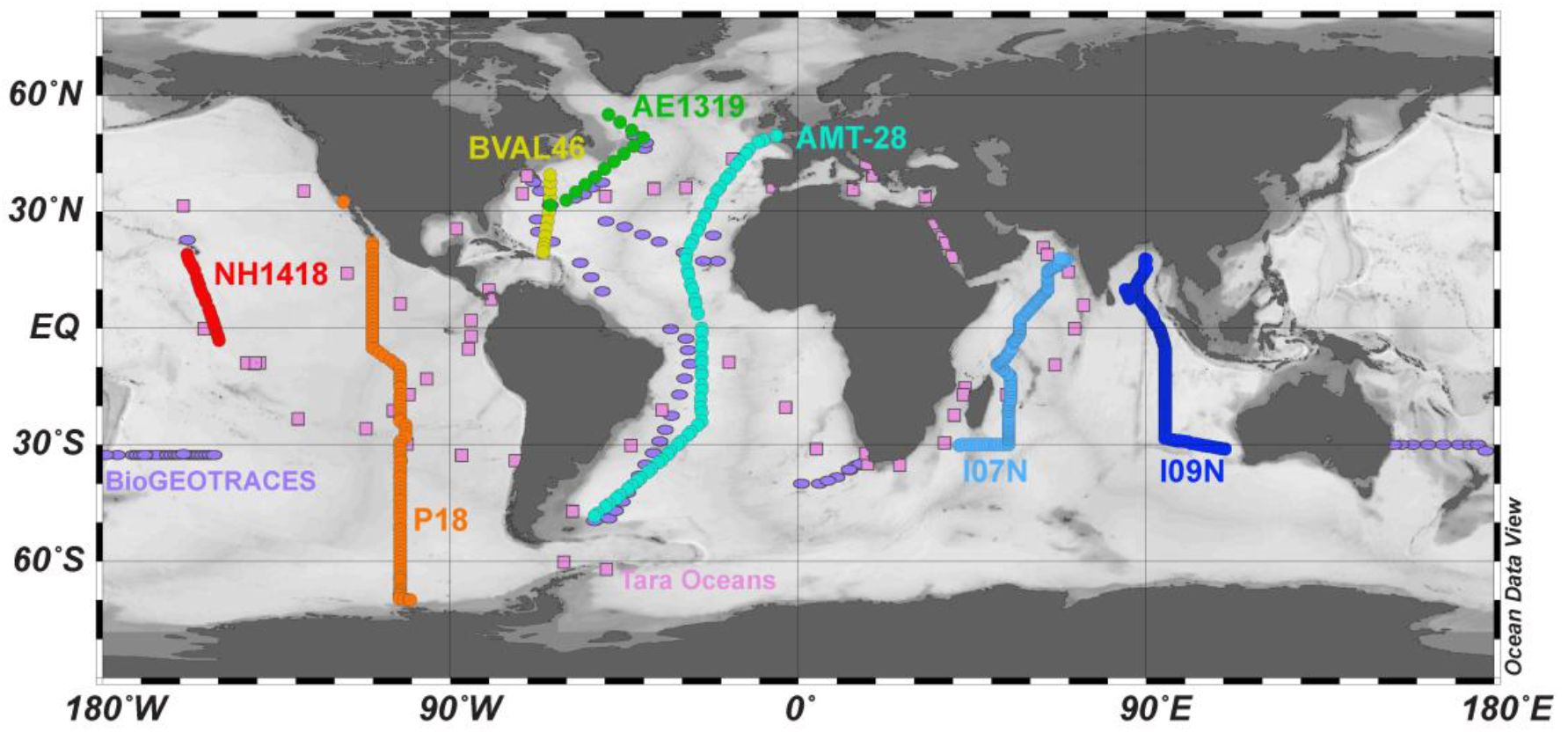
Distribution of global surface microbial metagenomes from Bio-GO-SHIP (circles) in comparison to Tara Oceans (squares) and bioGEOTRACES (ovals).

**Table 1:**
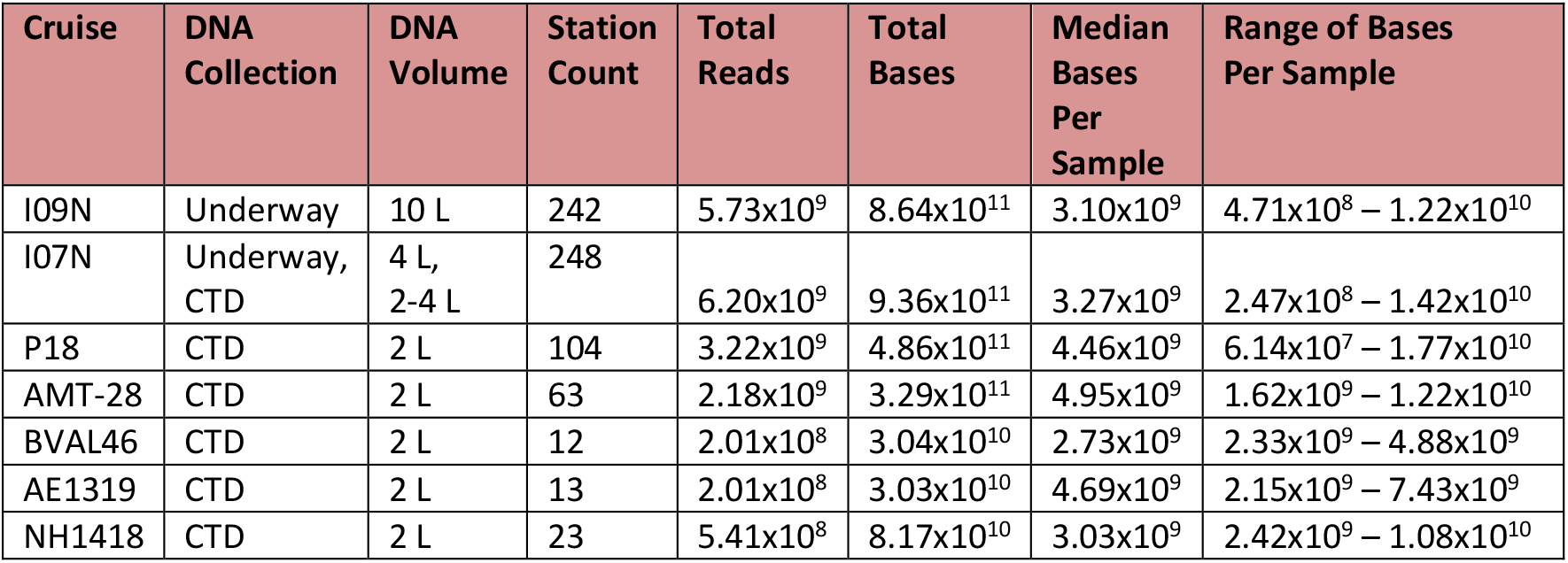
Sampling protocols and read counts for global Bio-GO-SHIP surface ocean metagenomes.

**Table 2:** Complete list of metadata variables collected on Bio-GO-SHIP cruises

| Campaign | Metadata Access | Metadata Variables |
| --- | --- | --- |
| I09N /<br>GO-SHIP | https://cchdo.ucsd.edu/cruise/33RR20160321 | Temperature, Dissolved O<sub>2</sub>, Conductivity, Density, Salinity, Nutrients (NO<sub>3</sub>, NO<sub>2</sub>, NH<sub>4</sub>, PO<sub>4</sub>, SiO<sub>4</sub>), Chlorofluorocarbons (CFCs) /SF<sub>6</sub>, Dissolved Inorganic Carbon (DIC), <sup>13</sup>C and <sup>14</sup>C of DIC, Total pH, Total Alkalinity, Stable gases (N<sub>2</sub>, N<sub>2</sub>O, Ar), <sup>18</sup>O, Chromophoric Dissolved Organic Matter (CDOM), Pigment HPLC, variable chlorophyll fluorescence, Dissolved Organic Carbon, underway Particulate Organic C N and P, underway pCO<sub>2</sub>, Lowered Acoustic Doppler Current Profiler, Chipods, Dissolved/ particulate/ cellular P and Fe, N P and Fe uptake rates, *Prochlorococcus*/ *Synechococcus*/ Picoeukaryotes/ Nanoeukaryotes cell counts |
| I07N /<br>GO-SHIP | https://cchdo.ucsd.edu/cruise/33RO20180423 | Temperature, Dissolved O<sub>2</sub>, Conductivity, Density, Salinity, Chlorophyll, Nutrients (NO<sub>3</sub>, NO<sub>2</sub>, PO<sub>4</sub>, SiO<sub>4</sub>), Dissolved Inorganic Carbon (DIC), Chlorofluorocarbons (CFCs) /SF<sub>6</sub>, <sup>14</sup>C of DIC, Dissolved Organic Carbon, Black Carbon, DO<sup>14</sup>C, Total pH, Total Alkalinity, Calcium, Dissolved Organics, Biomarkers, underway Particulate Organic C N and P, underway pCO<sub>2</sub> |
| P18 /<br>GO-SHIP | https://cchdo.ucsd.edu/cruise/33RO20161119 | Temperature, Dissolved O<sub>2</sub>, Conductivity, Density, Salinity, Nutrients (NO<sub>3</sub>, NO<sub>2</sub>, PO<sub>4</sub>, SiO<sub>4</sub>), Chlorofluorocarbons (CFCs) /N<sub>2</sub>O/SF<sub>6</sub>, Helium isotopes and noble gases (Ne, Ar, Kr, and Xe), Dissolved Inorganic Carbon (DIC), <sup>13</sup>C and <sup>14</sup>C of DIC, Total pH, Total Alkalinity, Stable gases (N<sub>2</sub>, O<sub>2</sub>, Ar), Dissolved Organic Carbon / Total Dissolved Nitrogen, Tritium, Black Carbon, DO<sup>14</sup>C/DO<sup>14</sup>C, Dissolved Organics, Biomarkers, underway Particulate Organic C N and P, underway pCO<sub>2</sub>, Wind speed, Wind direction, Air temperature |
| AMT-28 /<br>PML AMT<br>/ GO-SHIP | https://www.bodc.ac.uk/data/hosted_data_systems/amt/<br>https://amt-uk.org/Cruises/AMT28<br>https://cchdo.ucsd.edu/cruise/74JC20180923 | Temperature, Dissolved O<sub>2</sub>, Conductivity, Salinity, Nutrients (NO<sub>3</sub>, NO<sub>2</sub>, PO<sub>4</sub>, SiO<sub>4</sub>), biogenic silica/silicon uptake, Total pH, Total Alkalinity, Pigment HPLC, Chlorophyll a, Dissolved Organic C N and P, *Prochlorococcus*/ *Synechococcus*/ Picoeukaryote/ Nanoeukaryote/ Heterotrophic bacteria cell counts, FlowCAM, 15N/<sup>13</sup>C, Respiration (total, bacterial, size-fractionated), Bacterial production, underway Particulate Organic C N and P, underway wave radar (Cband), Sky Infrared Brightness temperature, Hyperspectral radiance/irradiance, Wind speed, Wind direction, Aerosol size/composition |
| BVAL46 /<br>NSF /<br>BATS | https://www.bco-dmo.org/project/2178 | Temperature, Dissolved O<sub>2</sub>, Conductivity, Salinity, PAR irradiance, Chlorophyll a, NO<sub>3</sub> + NO<sub>2</sub>, NO<sub>2</sub>, PO<sub>4</sub>, SiO<sub>4</sub>, Soluble Reactive Phosphorus (SRP), Particulate Organic C N and P, P uptake (max. uptake, half saturation conc.), *Prochlorococcus*/ *Synechococcus*/ Picoeukaryote/ Nanoeukaryote cell counts |
| AE1319 /<br>NSF | https://www.bco-dmo.org/project/2178 | Temperature, Dissolved O<sub>2</sub>, Conductivity, Salinity, PAR irradiance, Chlorophyll a, NO<sub>3</sub> + NO<sub>2</sub>, NO<sub>2</sub>, PO<sub>4</sub>, SiO<sub>4</sub>, Soluble Reactive Phosphorus (SRP), Particulate Organic C N and P, P uptake (max. uptake, half saturation conc.), *Prochlorococcus*/ *Synechococcus*/ Picoeukaryote/ Nanoeukaryote cell counts |
| NH1418 /<br>NSF | https://www.bco-dmo.org/project/2178 | Temperature, Dissolved O<sub>2</sub>, Conductivity, Salinity, Density, Fluorescence, PAR irradiance, Chlorophyll a, NO<sub>3</sub> + NO<sub>2</sub>, NO<sub>2</sub>, Soluble Reactive Phosphorus (SRP), Particulate Organic C N and P, *Prochlorococcus*/*Synechococcus*/ Picoeukaryote/ Nanoeukaryote / Crocosphaera cell counts |

**Figure 2:**
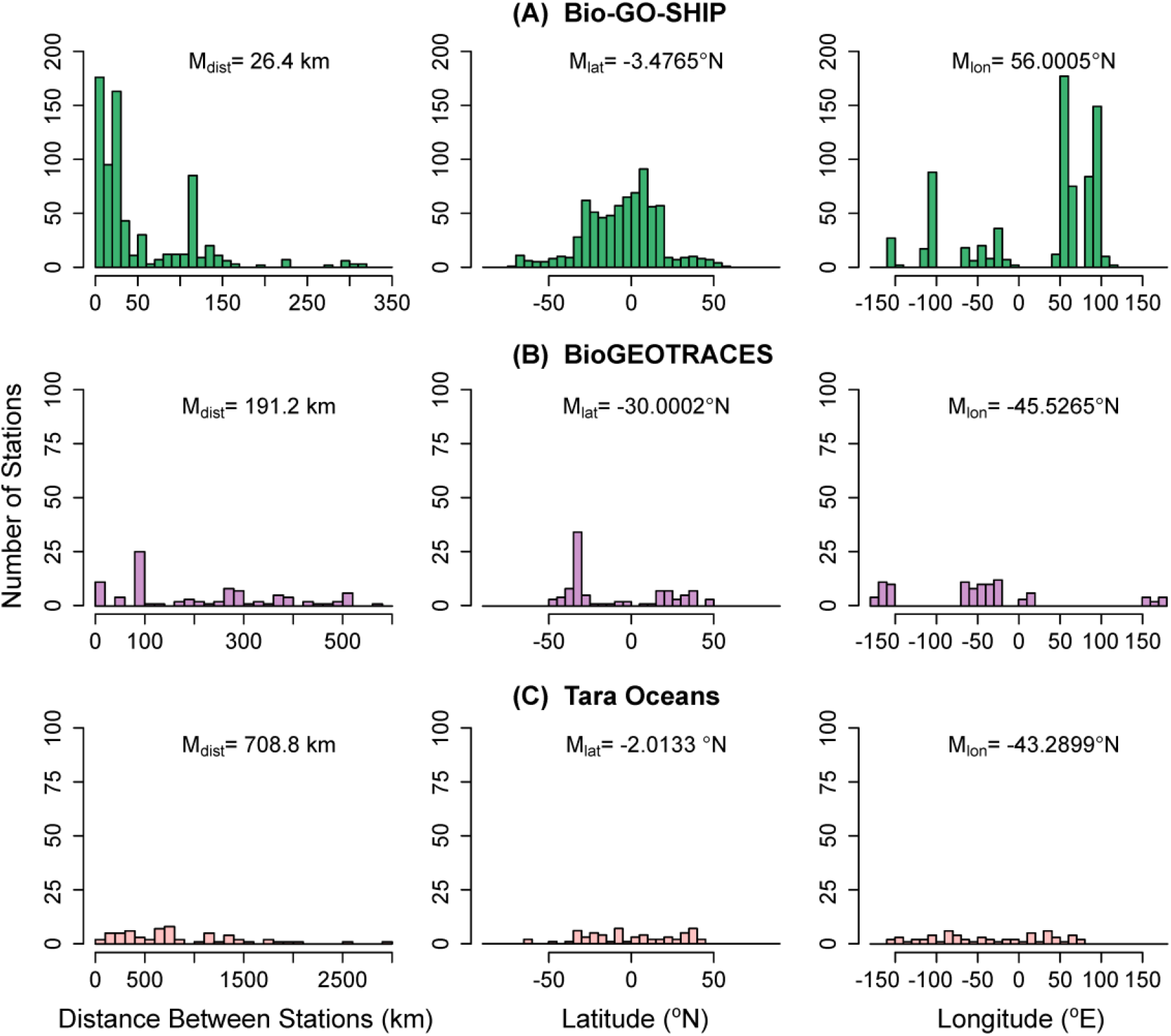
Comparison of the distance between stations, station latitudes, and station longitudes for global surface ocean metagenomes. Individual station locations from (A) Bio-GO-SHIP, (B) bioGEOTRACES and (C) Tara Oceans were examined. Plots are labelled with the median value, M. Station distance was calculated as the distance to the nearest station.

Due to their rapid generation times and high diversity, microbial genomes integrate the impact of environmental change^13^ and can be used a ‘biosensor’ of subtle biogeochemical regimes that cannot be identified from physical parameters alone^12, 14–16^. Thus, the fields of microbial ecology and oceanography would benefit from coordinated, high resolution measurements of marine ‘omics products (i.e., metagenomes, metatranscriptomes, metaproteomes, etc.). This dataset provides an important example of the benefits of a high spatial and temporal resolution sampling regime. Specifically, our data highlights the need for increased sampling of marine metagenomes in the Central and Western Pacific Ocean (Fig 1), areas above 50°N and 50°S (Fig 2), and below the euphotic zone. We hope and expect that these challenges will be addressed by the scientific community in the coming decade.

## Methods

On all cruises, whole (i.e., no size fractionation) surface water was collected via either the Niskin rosette system (depth ^~^3-5m) or the ship’s circulating seawater system (depth ^~^7m). Between 2-10 L of surface water (Table 1) was collected in triple-rinsed containers and gently filtered through a 0.22 μm pore size Sterivex filter (Millipore, Darmstadt, Germany) using sterilized tubing and a Masterflex peristaltic pump (Cole-Parmer, Vernon Hills, IL). DNA was preserved with 1620 μl of lysis buffer (4 mM NaCl, 750 μM sucrose, 50 mM Tris-HCl, 20 mM EDTA) and stored at −20°C before extraction.

To extract DNA (modified from Bostrom et al. 2004)^17^ Sterivex filters were incubated with 180 μl lysozyme (3.5 nM) at 37°C for 30 minutes followed by an overnight 55°C incubation with 180 μl Proteinase K (0.35 nM) and 100 μl 10% SDS buffer. DNA was extracted from the Sterivex with 1000 μl TE buffer (10 mM Tris-HCl, 1 mM EDTA), precipitated in an ice-cold solution of 500 μl isopropanol (100%) and 1980 μl sodium acetate (3 mM, pH 5.2), pelleted via centrifuge for 30 mins at 4°C, and resuspended in TE buffer in a 37°C water bath for 30 min. Next, DNA was purified using a genomic DNA Clean and Concentrator kit (Zymo Research Corp., Irvine, CA). Finally, DNA concentrations were quantified using a Qubit dsDNA HS Assay kit and Qubit fluorometer (ThermoFisher, Waltham, MA).

A total of 720 metagenomic libraries were prepared using Illumina-specific Nextera DNA transposase adapters and a Tagment DNA Enzyme and Buffer Kit (Illumina, San Diego, CA, cat. no. 20034197) (modified from Baym et al. 2015)^18–20^. Nextera adapter sequences to be used for bioinformatic quality trimming are: 5’-TCG TCG GCA GCG TCA GAT GTG TAT AAG AGA CAG-3’ and 5’-GTC TCG TGG GCT CGG AGA TGT GTA TAA GAG ACA G-3’. Custom Nextera DNA-style 8bp unique dual index (UDI) barcodes I7 (5’-CAA GCA GAA GAC GGC ATA CGA GAT [NNN NNN NN]G TCT CGT GGG CTC GG-3’) and I5 (5’-AAT GAT ACG GCG ACC ACC GAG ATC TAC AC[N NNN NNN N]TC GTC GGC AGC GTC-3’) were used to multiplex the metagenomic libraries. A total of 1 μl of 2 ng/μl DNA was added to 1.5 μl tagmentation reactions (1.25 μl TD buffer, 0.25 μl TDE1) and incubated at 55°C for 10 minutes. After tagmentation, product (2.5 μl) was immediately added to 22 μl reactions (1.02 μM per UDI barcode, 204 μM dNTPs, 0.0204 U Phusion High Fidelity DNA polymerase and 1.02X Phusion HF Buffer [ThermoFisher, Waltham, MA] final concentration). Barcodes were annealed to tagmented products using the following polymerase chain reaction (PCR): 72°C for 2 min., 98°C for 30 s., followed by 13 cycles of 98°C 10 s., 63°C 30 s., 72°C 30 s., and a final extension step of 72°C for 5 min.

To quality control tagmentation products, dimers that were less than 150 nucleotides long were removed using a buffered solution (1 M NaCl, 1 mM EDTA, 10 mM Tris-HCl, 44.4 M PEG-8000, 0.055% Tween-20 final concentration) of Sera-mag SpeedBeads (ThermoFisher, Waltham, MA). Metagenomic libraries were quantified using a Qubit dsDNA HS Assay kit (ThermoFisher, Waltham, MA) and a Synergy 2 Microplate Reader (BioTek, Winooski, VT). Libraries were then pooled at equimolar concentrations. Pooled library concentration was verified using a KAPA qPCR platform (Roche, Basel, Switzerland). Finally, dimer removal as well as read size distribution were checked using a 2100 Bioanalyzer high sensitivity DNA trace (Agilent, Santa Clara, CA).

54 samples were sequenced on two Illumina HiSeq 4000 lanes using 150 bp paired-end chemistry with 300 cycles (Illumina, San Diego, CA). All remaining samples were sequenced on three Illumina NovaSeq lanes using S4 150 bp paired-end chemistry with 300 cycles. The sequencing strategy produced a total of 1.83×10^10^ reads, or 2.75×10^12^ bp. The median number of bases per sample was 3.48 billion (range: 61,400,000 – 17.7 billion). The sequencing cost per bp in US dollars was $8.2×10^−9^.

## Data Records

The majority of the samples here were collected under the auspices of the international GO-SHIP program and the national programs that contribute to it^21–24^. A comprehensive data directory of metadata resources is available at https://www.go-ship.org/. Bottle data and cruise report links are provided in Table 1.

Metadata variables from the AMT-28 cruise (https://www.amt-uk.org/) are hosted by the British Oceanographic Data Centre, and may be requested through the following URL: https://www.bodc.ac.uk/. Select metadata are also available through GO-SHIP^24^.

The BVAL46, AE1319, and NH1418 cruises were collected as a part of the “Biological Controls on the Ocean C:N:P Ratios” project funded by the NSF Division of Ocean Sciences^25–28^. Data associated with these deployments are hosted by the NSF Biological and Chemical Oceanography Data Management Office (BCO-DMO). A comprehensive list of metadata resources is available at https://www.bco-dmo.org/project/2178.

All sequencing products associated with the Bio-GO-SHIP program can be found under BioProject ID PRJNA656268 hosted by the National Center for Biotechnology Information Sequence Read Archive (SRA)^29^. SRA accession numbers associated with each metagenome file are provided in Supplementary Table 1.

## Technical Validation

To ensure that no contamination of metagenomes occurred, negative controls were used. To ensure optimum paired-end short read sequencing, a 2100 Bioanalyzer high sensitivity DNA trace (Agilent, Santa Clara, CA) was used for each library to confirm that 90% of the sequence fragments were above 250 bp and below 600 bp in length. Qubit (ThermoFisher, Waltham, MA) and a KAPA qPCR platform (Roche, Basel, Switzerland) were used to ensure that all pooled libraries were submitted for sequencing at a concentration >15 nM.

## Usage Notes

The genomic data described here have not been pre-screened or processed in any way. We recommend quality control parameters. Prior to our sequence analysis in subsequent projects, we removed adapter sequences, performed sequence quality control, and ensured there was no contamination from common genomic add-ins such as Phi-X using the following code parameters:

Trimmomatic (v0.35): PE ILLUMINACLIP:NexteraPE-PE.fa:2:30:10 SLIDINGWINDOW:4:15 MINLEN:36
BBMap (v37.50): bbduk.sh -Xmx1g ref=/BBMap/37.50/resources/phix174_ill.ref.fa.gz k=31 hdist=1

## Code Availability

Custom scripts were not used to generate or process this dataset. Software versions and non-default parameters used have been appropriately specified where required.

## Acknowledgements

We would like to thank the captains and crew of the R/V Atlantic Explorer, R/V New Horizon, R/V Ronald H. Brown, R/V Roger Revelle, and the R.R.S. James Clark Ross. Financial support for this project was provided by the National Science Foundation (OCE-1046297, 1559002, 1848576, and 1948842 to ACM). LB and DLV were supported in part under the auspices of the Cooperative Institute for Marine and Atmospheric Studies (CIMAS), a cooperative institute of the University of Miami and NOAA (cooperative agreement NA10OAR4320143). The PML AMT is funded by the UK Natural Environment Research Council through its National Capability Long-term Single Centre Science Programme, Climate Linked Atlantic Sector Science (grant number NE/R015953/1). This study contributes to the international IMBeR project and is contribution number ### of the AMT program. GO-SHIP is supported by NOAA Global Ocean Monitoring and Observation program (U8R1SE3-PRF) and by the National Science Foundation (OCE-1437015). P18 and I07N were NOAA-led cruises and I09N was an NSF-led cruise.

## Author contributions

A.A.L. wrote the manuscript, coordinated sample collection, collected/processed samples, designed protocols, performed metagenomic sequencing, and compiled metadata.

C.A.G. coordinated sample collection, collected/processed samples, performed metagenomic sequencing, and compiled metadata.

M.L.B performed metagenomic sequencing.

J.A.L. collected/processed samples and performed metagenomic sequencing.

N.G. coordinated sample collection and collected samples.

L.J.U. processed samples and compiled metadata.

L.B, B.G.C., R.E.S., L.T., and D.L.V. coordinated GO-SHIP collection and collaboration efforts.

G.T. coordinated PML AMT/GO-SHIP collection and collaboration efforts.

A.C.M. designed and supervised the study, secured funding, and coordinated GO-SHIP collection.

All authors contributed to manuscript editing and revision.

